# Spatiotemporal analysis of speckle dynamics to track invisible needle in ultrasound sequences using Convolutional Neural Networks

**DOI:** 10.1101/2022.08.02.502579

**Authors:** Amin Amiri Tehrani Zade, Maryam Jalili Aziz, Hossein Majedi, Alireza Mirbagheri, Alireza Ahmadian

## Abstract

**Objective:** Accurate needle placement to the target point is critical for ultrasound interventions like biopsies and epidural injections. However, aligning the needle to the thin plane of the transducer is a challenging issue as it leads to the decay of visibility by the naked eye. Therefore, we have developed a CNN-based framework to track the needle using the spatiotemporal features of speckle dynamics.

**Methods:** There are three key techniques to optimize the network for our application. First, we proposed a motion field estimation network (RMF) to extract spatiotemporal features from the stack of consecutive frames. We also designed an efficient network based on the state-of-the-art Yolo framework (nYolo). Lastly, the Assisted Excitation (AE) module was added at the neck of the network to handle imbalance problem.

**Results:** Ten freehand ultrasound sequences are collected by inserting an injection needle steeply into the Ultrasound Compatible Lumbar Epidural Simulator and Femoral Vascular Access Ezono test phantoms. We divided the dataset into two sub-categories. In the second category, in which the situation is more challenging and the needle is totally invisible statically, the angle and tip localization error were 2.43±1.14° and 2.3±1.76 mm using Yolov3+RMF+AE and 2.08±1.18° and 2.12±1.43 mm using nYolo+RMF+AE.

**Conclusion and significance:** The proposed method has the potential to track the needle in a more reliable operation compared to other state-of-the-art methods and can accurately localize it in 2D B-mode US images in real-time, allowing it to be used in in current ultrasound intervention procedures.

## I. Introduction

**R**eal-time and accurate placement of the hand-held needle is vital for many medical procedures including epidural anesthesia, biopsies and, treatment injections. These procedures are being guided using manual palpation or imaging techniques such as c-arm and fluoroscopy. Recently, Ultrasound (US) have gained more attention in image-guided interventions due to its portability, real-time operation, and non-ionizing imaging [1], [2]. However, due to the specular reflection of the needle surface, it appears as ambiguous, discrete line-like features in US images. Also, from the operator’s viewpoint, aligning the needle with the transducer’s narrow field of view is a difficult task. This difficulty is further exacerbated by poor visible reflection. To alleviate these challenges, two main approaches exist, including hardware-based [3]– [10] and image-based approaches. The main drawback of using peripheral devices like mechanical needle guides, embedded sensors and robotics [11], are the limitations and the need for additional equipment, which are certainly not part of the current clinical practice.

In image-based methods, the US frames are processed statically or dynamically. In static analysis, the frames are investigated independently of time, assuming that the needle shaft is seen as a line-like brighter intensity in the images. There are several traditional methods to extract lines in US frames using statistical analysis of image intensity [12], hough transform [13]–[16], projections[17]–[21], random sample consensus(RANSAC) [22]–[24][25], filtering[10], [26]–[28] and radon transform[29], [30]. Log-Gabor filters have also been used to extract phase images of needle projections. It shows that the phase is more robust to intensity variation than amplitude for segmenting bone and the needle surface in US images [12].

Another strategy to extract needles in US-based navigation systems is to process frames time-dependently. The prominent discriminative features appeared as dynamic varying of speckles in these dynamic approaches. The invisible periodic pattern of US features was examined in a dynamic method [31]. This method demonstrated the effectiveness of utilizing spectral properties of the flow filed to detect macro blocks that have the maximum correlation with the reference tremor pattern [31]. Compared to traditional approaches, machine learning has the potential to incorporate prior knowledge into inference. Continuing the previous work, a study was conducted based on volumetric features derived from the spatiotemporal analysis of the motion field phase. In this method, phase images were estimated using the steerable pyramid of multiple oriented Gabor filters. The spatiotemporal and spectral features extracted from the raw and differential flow field were used to train SVM to classify each pixel into needle or background regions [32]. Additionally, some studies used statistical methods to model the phase variation of pixel intensities. After estimating pixel-wise phase features by applying three oriented Gabor filters in two scales, temporal features are extracted from the resulting multiscale images to model an Auto-Regressive Moving-Average (ARMA) system. Finally, SVM was used to classify each pixel and Hough transform to estimate line parameters from the probability map [33].

In recent years, many studies have been conducted toward using Convolutional Neural Networks (CNN) in medical image segmentation [34] and specifically US-guided needle detection applications. The first use of the CNN to track the needle shaft and tip in US images was performed statically. It used the Regional Proposal Network (RPN) and fast R-CNN as a state-of-the-art object detection network to detect the region of interest of the needle shaft and then classify them [35]. Next, to handle low intensity appearance of the needle, basic logical operation was used to enhance the moving needle in a dynamic background [36]. CNN-based approaches have the potential to extract more discriminative dynamic features, and therefore, in recent years, many studies in the field of US-guided needle detection have focused on the use of CNNs. A modified version of the U-Net was proposed to segment needle-like interventional instruments in 2D US images [37]. They tested their proposed network on several clinical procedures, including gynecologic, brachytherapy, liver ablation, kidney biopsy and ablation, and phantom studies. A two channel encoder U-Net was introduced to capture needle motion from two sequential frames. It includes a coarse to fine framework, in which the extracted ROIs from the adjacent frames are fed to the U-Net [38]. In all of these methods, it is assumed that the intensity in the vicinity of the needle is higher than in the background and can therefore be detected using the intensity threshold or phase of the image. This hypothesis is valid if the needle is entirely in-line with the transducer plane and is sufficiently perpendicular to the incident beam. Unfortunately, so many of these methods are failed in clinical usage. Therefore, estimating the motion model from the sequential frames is the target of several studies on US-guided needle tracking methods. Lucas-Kanade optical flow was used to track the selected cannula segment edges in the US plane. Furthermore, the out-of-plane pattern of the intersection cannula is detected using circular Hough transform [13]. By concatenating Long Short Term Memory (LSTM) and CNN blocks, a deep learning framework has been proposed to model the temporal properties of the tip in US sequences [39]. The input is made by integrating raw and the enhanced US frames. To enhance the needle tip, it used bitwise logical operation and median filter [39]. In this study, where LSTM was used to track spatial features in time, speckle dynamics were never considered as a discriminative feature. Also, an affine motion model was proposed to track the needle tip in its vicinity. It showed that the thin-plate spline (TPS) is superior to the affine model for modeling tissue deformations [10].

Some gaps are ignored in the studies to segment or detect needles in US frames. Firstly, almost all the reviewed studies are based on independently tracking the spatial features extracted from the frames. Whereas the reflection of the incident wave at various angles is added in time and leads to a random vibration of high-intensity speckles. These vibrating speckles have a discriminative pattern over time as a one-dimensional signal and can be extracted using dense motion field estimation. Secondly, the main problem in object detection frameworks is their poor capability to confront class imbalances and hard negative examples which severely degrades the accuracy of predictions.

We used Motion field estimation methods to utilize the speckle motions in the tracking needle framework to resolve the first issue. Motion field estimation has been studied in various applications. In traditional optical flow like Farenback’s method [40], displacement field is measured by solving under-determined equations using the intensity gradient of pixel. On the other hand, motion field estimation can be obtained by measuring a flow field that aligns the consecutive frames. Recently, many studies have used supervised deep learning methods to estimate flow in which the flow field is required for each image pair as a ground truth [41], [42]. These networks include both a CNN and spatial transformation function [43] to wrap the moving frame using the predicted flow field. An unsupervised end-to-end network has been proposed to estimate the deformation between two pairs of images. This study has two main advantage which suit our approach. First, this technique does not confront flow field gathering as ground truth or synthetic dataset creation [42] for training as a fundamental problem in supervised motion field estimation. Secondly, this network is worked in one-stage end-to-end fashion that could be embedded as an independent box to current needle detection networks [44]. Considering these advantages, we augmented our network with this method to estimate the motion filed between two consecutive US frames. Augmenting CNNs with other traditional optical flow methods or CNN-based motion field networks and sensors to measure the speckle’s motions has been shown to improve its predictive capabilities in US practices [45].

The attention mechanism was utilized to improve the training for the second problem. Attention was studied in various frameworks. Lin et al [46] introduced the “focal loss” to handle the foreground-background imbalance problem. By utilizing this approach, the network learns to focus more on hard negative examples. Some studies take a different approach to solve the problem. Derakhshani et al [47] used ground truth bounding boxes to excite more effective activations during training. The activations in the vicinity of the objects are excited more at the beginning of the training and gradually decreased to zero until the end of training. This technique is based on Curriculum Learning (CL), which has shown to be effective in convergence speed and the final accuracy [48].

This paper presents a new image guidance method to track invisible needles in US-guided interventions. According to the proposed method, the stack of US images is combined with displacement map images to augment training with movement features. We used traditional optical flow and CNN-based motion field estimation models to extract spatiotemporal properties of the speckle dynamics. These augmented images are then fed to the CNN-based needle detection network. During training prominent activations are spatially excited to attend more on the object neighborhood rather than the background. The proposed algorithm was trained and tested on US videos captured using epidural and femoral vascular access phantoms. To the best of our knowledge this is the first work that studies the effect of speckle dynamics in one-stage end-to-end needle detection networks. Furthermore, the attention mechanism was utilized as a CL paradigm to improve the unbalancing between the needle and the background regions.

## II. Materials and methods

The proposed network is designed to detect needles based on the spatiotemporal properties of the speckle dynamics. Therefore, to achieve these goals, we utilized three CNN-based techniques, which are,

- Motion field estimation
- Assisted Excitation (AE)
- Needle shaft and tip detection

These methods are studied in order to develop a clinically acceptable system with the following characteristics,

- Consistent operation in real-world situations where the needle appears partially or with a minimum manifestation of intensity, in which the distinguishing features emerged as a low frequency speckle dynamic.
- Achieve an end-to-end network with real-time operation
- Compatible with current clinical setup

The following sections express the designed motion estimation network and compare the traditional motion estimation methods based on the optical flow. Afterward in section *B* the attention mechanism and, finally, the main detection network is explained.

### A. Motion Field Estimation (MFE)

To capture spatiotemporal features, we selected and compared a traditional optical flow and a CNN-based unsupervised deformable field estimation method. Gunnar-Farneback [40] (GF) was employed as a traditional dense motion field estimation method. This approach calculates the field by minimizing the least square error in an under-determined problem. The brightness constancy constraint is the basic assumption that degrades the efficiency of US-based systems. This assumption is not always fair because the reflections appear as moving speckles in invisible needle detection frameworks. Many studies utilize CNNs to measure optical flow. In these approaches, each pair of consecutive frames is fed to the CNN, and the predicted tensor is compared with the true field as a ground truth. The key drawback of these techniques is their ground truth, which is often absent in many settings. FlowNet [41], FlowNet2[49] and SpyNet [42] are among the popular CNN-based networks in this field.

Using another approach, dense correspondence is estimated by employing deformable registration algorithms to solve a constrained optimization problem. Despite the success of traditional methods [50], [51], CNN-based registration procedures have recently attracted more attention. Besides the online capacity of these frameworks, they are more precise and robust to intensity and structural fluctuations. Some strategies made use of Spatial Transformer [43] for registration., Training requires a pair of images or volumes within these approaches, and the loss is measured between the fixed and the wrapped images using Spatial Transformer block. This paper used a modified version of VoxelMorph [44] to predict the motion field. This network could be included into the needle detection network efficiently and in an end-to-end manner.

Our methodology aims to capture speckle motion patterns at multiple scales and make pixel-wise predictions, so losing the resolution due to max-pooling and down-sampling could not be the best option. Instead of using the encoder-decoder structure, we employed resolution-preserving dilated filters to extract deep features without sacrificing the details. This technique has been studied in object detection [52] and semantic segmentation [53] applications. As shown in Fig. 1, the stacked images are processed using 5 dilated convolution filters at rates of 1, 2, 4, 8, and 16.

**Fig. 1.**
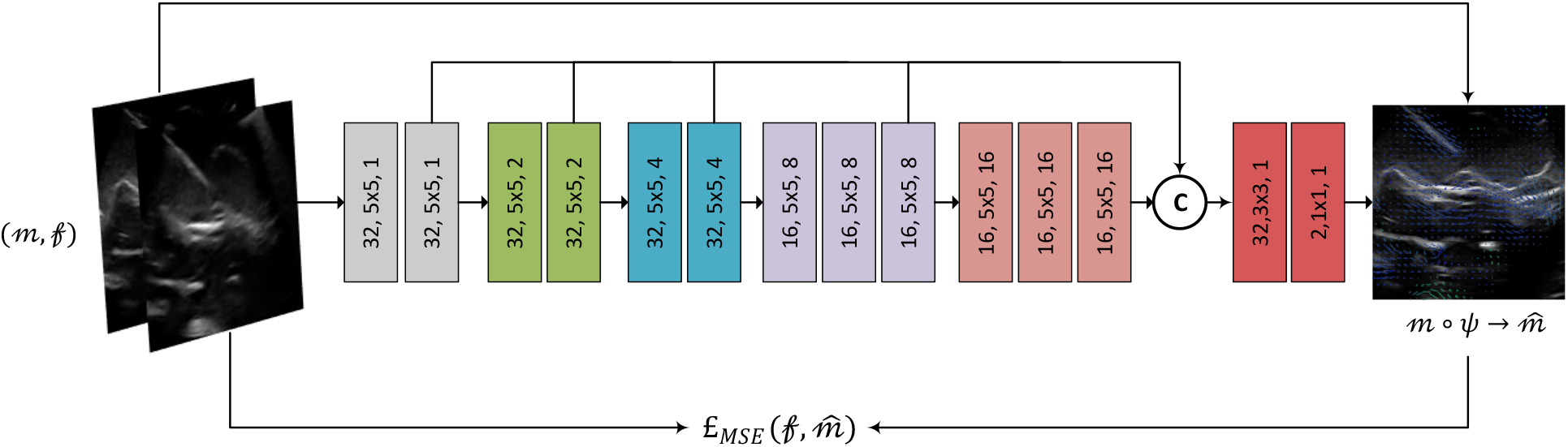
Motion field estimation network. Multiscale features are captured using dilated convolutions. The output tensors from 5 scales are concatenated and finally the flow field is estimated using two 1D convolutional filters. £_MSE is the measure of similarity between the moved and the fixed frames.

The output tensors on each scale are aggregated for processing in the subsequent stages. Finally, the motion field is predicted using two 1D convolutions. The anticipated field is applied to the moving frame and the loss is measured as a criterion of similarity between the moved and the fixed frame. We named it the Resolution-preserving Motion Field (RMF) network.

Let 𝒻 ∈ ℝ^*w*×*h*^, *m* ∈ ℝ^*w*×*h*^ and 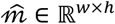 be the fixed, moving and the moved images respectively. As seen in (1), the moved image is obtained by transforming *m* using the predicted field (*ψ*).

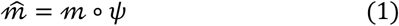

We utilized Mean Square Error (MSE) to measure the loss function. This function could be measured using (2).

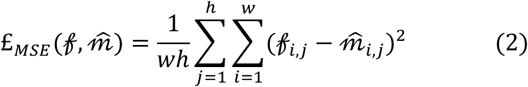

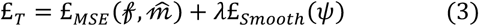

We also used £_*Smooth*_ to regularize the network into more smooth variations in *ψ*,

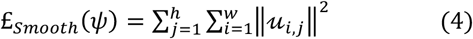

In (3), *λ* is the regularization parameter. The motion field gradients (*u*) are employed in (4) to minimize the overall variations in the filed vectors.

### B. Attention mechanism

One-stage detectors have the shortcoming of predicting plenty of bounding boxes, many of which are negative examples. The classification of these negative examples is simple, making them vulnerable to biased training. Furthermore, the class imbalance is a significant issue in many cases where the foreground is trivial compared to the background. Therefore, it is helpful to train the network with important areas by attending more to foreground regions. Assisted excitation (AE) of activation is a strategy to enhance recall by employing ground truth bounding boxes in the training process. The proposed attention in the AE structure was adapted from Derakhshani et al. [47] and is shown in Figure 2a. The excitation was done by applying the ground truth bounding boxes on the filtered tensor. We utilized one 1D convolution filter instead of “average over channels” block to capture features by trainable weights. This technique could be more beneficial than “average over channels” because there are many irrelevant activations over channels that could degrade the filtering response. The network must learn the prediction without using the ground truth since the bounding boxes are unavailable during the inference stage. The CL approach was utilized to handle the attention in a semi-supervised fashion. Using this technique, the excitation factor (attention coefficient) was set to one at the start and gradually decreased to zero until the end of training.

**Fig. 2.**
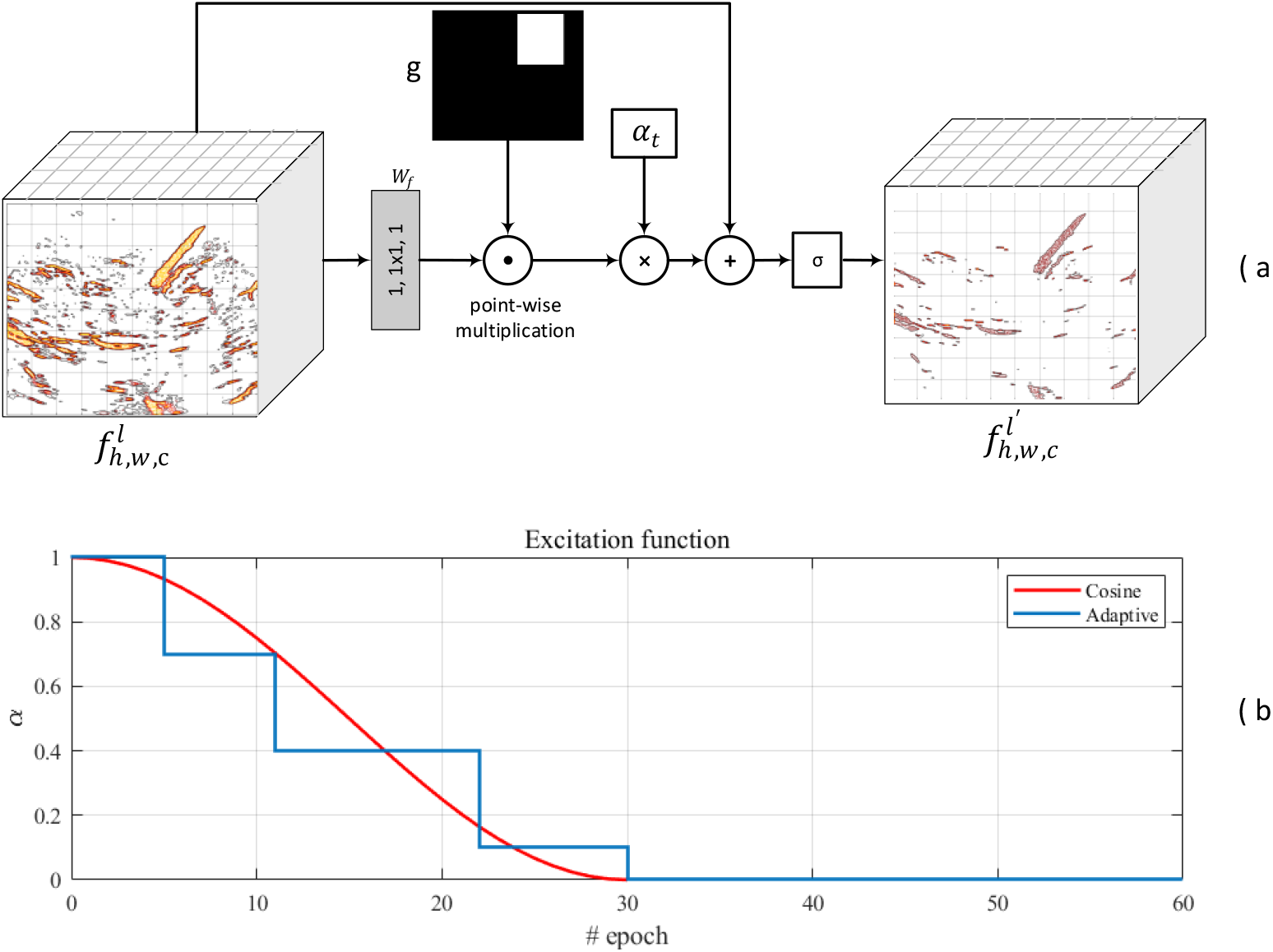
Attention mechanism, a) considering (5–7) the attended feature map 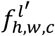 is obtained by applying the excitation factor (*α*) in a decreasing rate. The overall structure is similar to the work presented by Derakhshani et al[47] except for the *W*_*f*_ and the adaptive excitation factor, b) adaptive excitation factor profile. Using this technique, the excitation factor is reduced when the metric does not improve on validation.

Let 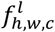 and 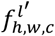 be the feature maps before and after applying the AE in layer l. We employed one 1D convolution filter (*W*_*f*_ ∈ ℝ^*c*×1^) to map the activations to the R^1 dimensional tensor. The filtered tensor is masked by g, the ground truth bounding box. As seen in (6) and (7), the resulted tensor is derived by applying the sigmoid activation function (σ) to the excited activation map 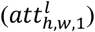. Fig. 2.a represents a schematic of the AE module.

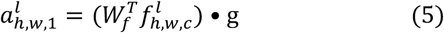

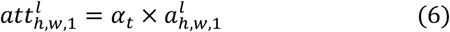

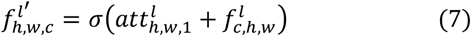

We proposed an adaptive version of the excitation factor instead of a fixed monotonic function [47]. Rather than employing a smooth, monotonically decreasing function (8) to alter the attention coefficient, we degraded the parameter stepwise, after a few epochs with no enhancement in validation accuracy. This strategy improves the network’s convergence as well as its overall performance. The adaptive excitation function is depicted in Fig. 2b.

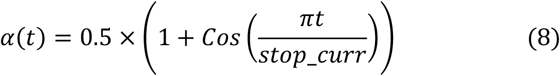

### C. nYolo, needle detection network

Across CNN-based object detectors [54–57], Yolo has been among the most successful single-stage end-to-end ones. This technology has higher real-time capabilities than other two-stage object detectors. Nevertheless, these networks are more susceptible to false positive predictions. Yolo framework is released in three versions, Yolov1[58], Yolov2[59] and Yolov3[55]. Among these, Yolov3 was the last idea of Redmon in which the Feature Pyramid Network (FPN) [60] was used to extract predicted boxes on multiple scales. Yolov4[56], Yolov5[61] and PP-Yolo [57] are Yolo’s most recent versions, which were published after 2020. Yolov4’s main contribution was to change the training strategy without lowering the inference efficiency. They improved data augmentation, semantic distribution bias, and bounding box regression objective function as part of the Bag of Freebies paradigm. Besides the overlaid methods on the training process, the Bag of Specials enhances the detection accuracy against decreasing the inference efficiency. Furthermore, Yolov4 used PANet [62] instead of FPN for multiscale processing. The overall architecture of Yolov5 is same as Yolov4 but with some difference in data augmentation and learning process and also 90 percent parameters decrease and so faster inference. Another approach to improve the efficiency of Yolov3 learning was PP-Yolo. The backbone and head of Yolov3 was used as main building blocks, and only the learning process was improved to achieve balance between effectiveness and efficiency. Another research was done to add attention to the training process. Assisted excitation of activations was proposed to use the bounding box masks as an attention module to add more gain on target areas. The proposed idea was shown to improve the accuracy with less false positive rate [47].

We presented a network based on Yolov3 to recognize the needle and the relative tip more efficiently. Some enhancements are offered to adapt the network in our case,

- Residual block has been found to have good performance based on accuracy and convergence rate in many object detectors, including Yolov3. Therefore, we utilized it as the main building block in our experiment to route each feature map directly to the next layer.
- Speckle motion patterns are detectable spatially in the beginning stages of the network but deeper in temporal features. So first the motion features are extracted by feeding the stack of US frames to the RMF block and extracted spatiotemporal feature tensors are added elementwise and routed to the input of the detection network backbone.
- We employed Atrous Spatial Pyramid Pooling (ASPP) [63] to extract deep features while maintaining the resolution in the last stage of the backbone. It is basically a more receptive field needed for the shaft than the tip.
- We benefited from the AE module in training. Using this technique, the network learns to pay more attention to target areas. Therefore, aside from ignoring easy negative examples and handling the class imbalance problem, the accuracy of the prediction has improved in terms of *Sensitivity* and *Specificity*.

The overall structure of the proposed network consists of two sections, Spatiotemporal Feature Extractor and Detector. In the first section, the motion field’s amplitude and phase of successive frames are recognized using the RMF module. Afterward, these stacked frames and the associated amplitude and phase images are passed through two Residual Blocks with 32 convolutional filters to extract spatiotemporal features and sent them to the Detector section. As shown in Fig. 3, after 3 levels of downsampling in the backbone, the resulting feature maps in the last layer is generated by utilizing the ASPP block. The generated feature maps in each layer of the encoder are extracted using two one dimensional convolution filters of sizes 32 and 6 to predict pixel-wise line parameters and the confidence score of the needle and the corresponding tip. The output prediction vector of size 6 consists of three normalized line parameters (*t*_*x*_, *t*_*y*_, *θ*), abjectness score *c*^*n*^ and two confidence scores for the needle shaft and tip (*c*^*l*^, *c*^*t*^). Although the AE module can be included at any stage, we experiment in multiple places and get the best results by adding it at the beginning of the max-pooling operation in the neck portion.

**Fig. 3.**
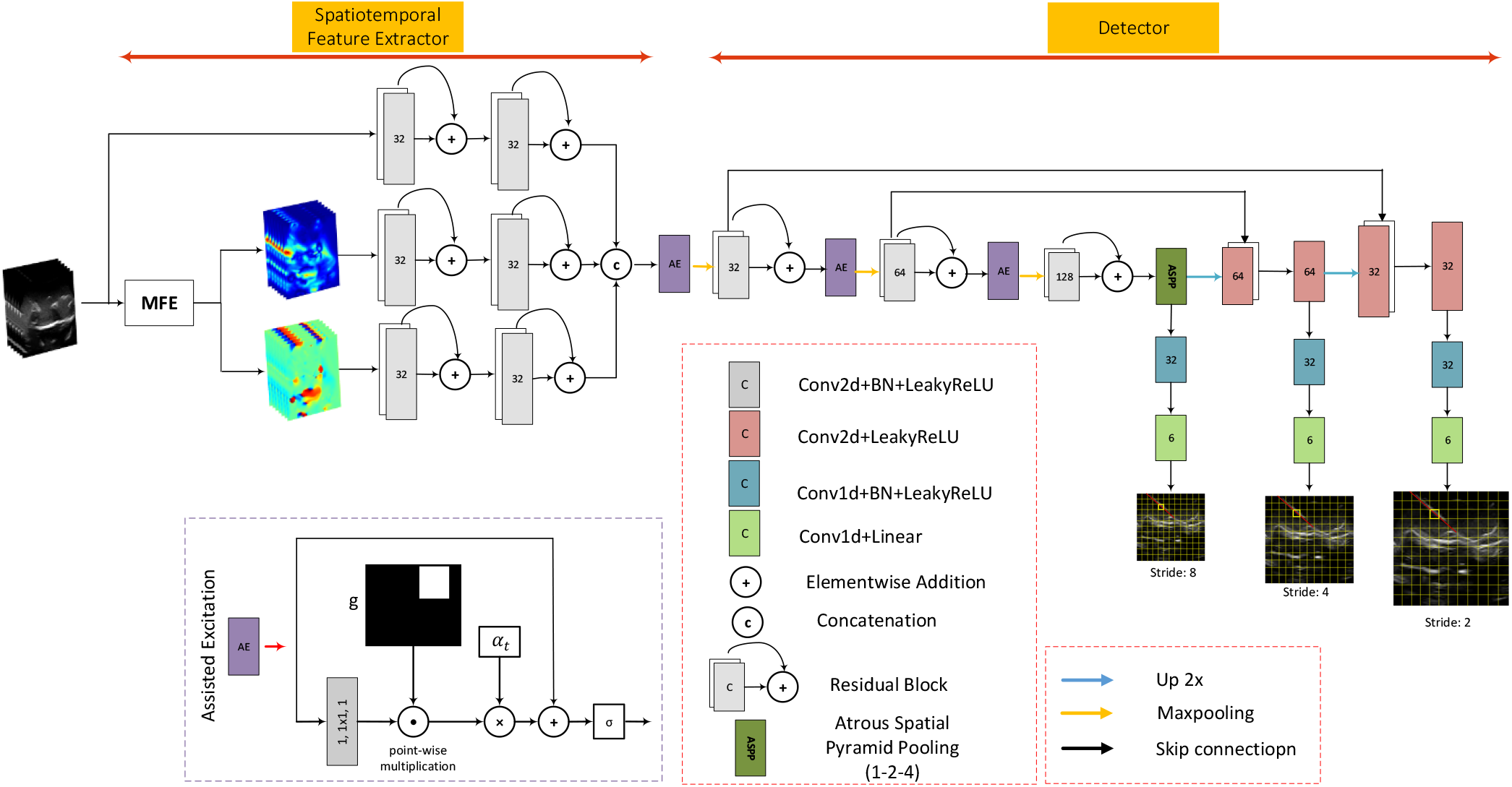
nYolo architecture. The proposed network is designed based on Yolov3 with residual blocks as an efficient feature extractor in the encoder section. Skip connections were utilized to recover the resolution in the neck using FPN block. Finally, the dense prediction framework obtains the output tensor in three scales. The speckle movement dynamic is retrieved using the motion field estimation network (MFE). AE modules are added before each max-pooling.

We used a multi-part loss function as in basic Yolo [58] to train the nYolo network. As shown in (9), the total loss is composed of three parts, the Needle-ness loss (£_*N*_), the confidence score of the existence of the needle (£_*C*_) and the regressor loss (£_*L*_) to predict line parameters in each grid cell. The term 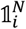 denotes the existence and 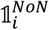 the absence of the needle in cell i. We could direct the training toward penalizing misclassification in cells with the needle by weighting the loss as in (10). We chose 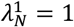 and 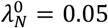to handle this imbalance between foreground and background. In (12) and (13), the line parameters are normalized between 0 and 1. So the Euclidean distance of the line and cell center is divided by the image’s diagonal size in the corresponding scale (d) and θ is divided by π/2.

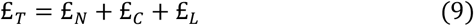

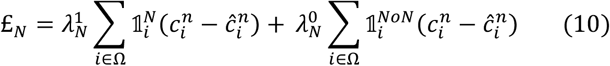

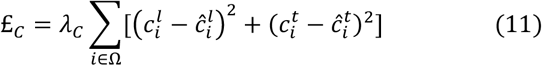

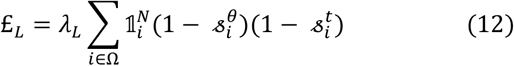

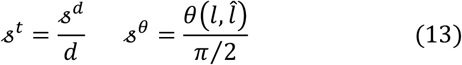

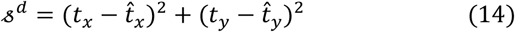

We used EA-score to penalize the line parameter discrepancy as in (12). In this metric, the Euclidean and the angular distances are considered for computing the loss function. It is shown that this metric has more consistent results than IOU-based line discrepancy measurement metrics [64]. As seen in Fig. 4, the Euclidean distance is measured between the center of two lines in the current grid cell.

**Fig. 4.**
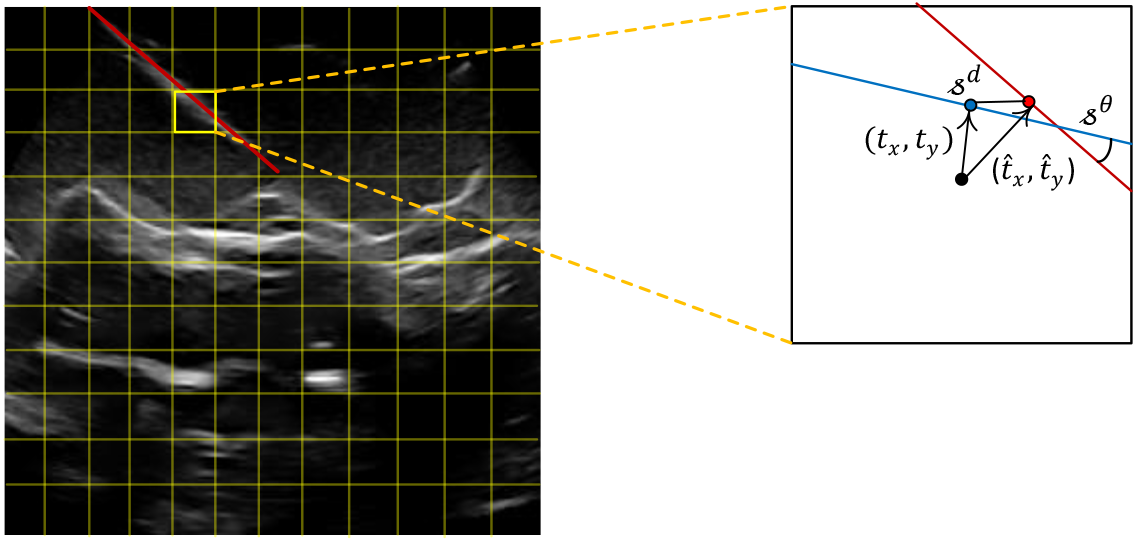
Evaluation metric between the ground truth and the predicted line parameters. These measurements are done on three scales. As seen in (13) and (14), 𝓈^*d*^ and 𝓈^*θ*^ are the metrics to measure the distance between the ground truth and the predicted line parameters. (*t*_*x*_, *t*_*y*_) and 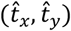 are the offsets from the cell center in the ground truth and the corresponding predicted line parameters.

### D. Inference

The output tensor has a per-cell prediction on three scales. In contrast to earlier versions of object detection systems, in our application there is only one needle and tip in one frame. This limitation was utilized to determine the optimum line and its associated tip. An example of the predicted abjectness score (*c*^*n*^) in three scales of the output layers are depicted in Fig. 5. The predictions are more consistent in term of *Sensitivity* (15) and *Specificity* (16) in a lower resolution output and, therefore, more reliable than those in the higher resolution. There are some outliers as False Positive predictions, which severely degrades the final detection accuracy in terms of Specificity. Higher resolution predictions are more prone to false alarms than lower resolution predictions, but they are more accurate in EA-score. So we designed a hierarchical detection algorithm that start from the lower resolution and gets more accurate by following into higher resolution predictions. Lower resolution predictions are used to select the line candidates and get enhanced by averaging the matched ones in the following steps.

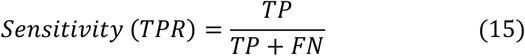

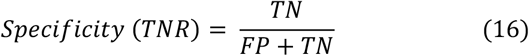

**Fig. 5.**
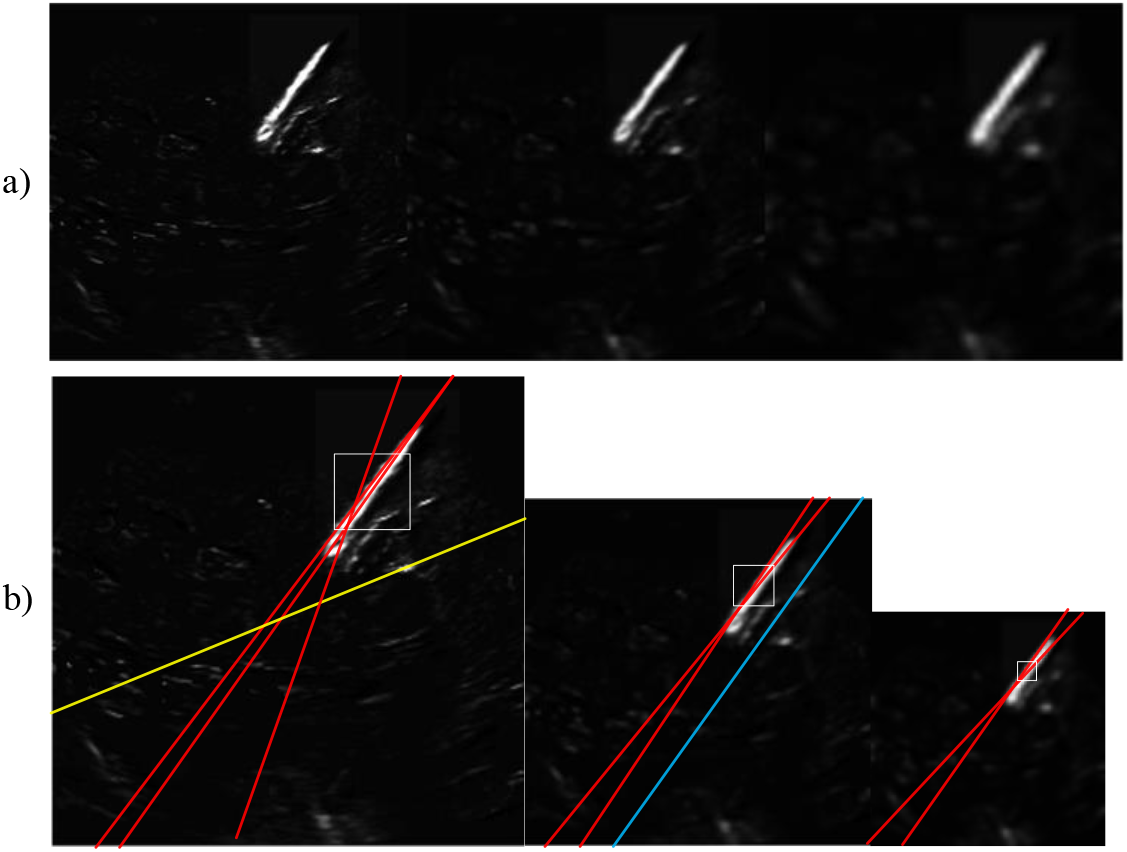
a)The predicted needle confidence score in three scales 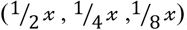, b) Corresponding line candidates in tree scales to be processed in line detection stage. The output tensor gets per-pixel line parameters with confidence scores. These candidates are processed to calculate the best which matches the predicted lines. As be seen, the predicted lines in lower resolution have high accuracy but in the price of more vulnerable to outlier (yellow lines)

Let 𝒯^2^, 𝒯^4^ and 𝒯^8^be the predicted tensors in three scales. Per-cell prediction vectors corresponded to the offset from the grid cell center (m), the angle (θ) and the confidence score of the needle shaft and tip respectively (*c*^*l*^, *c*^*t*^). In our experience, more robust and outlier-free detection was carried out in the lower scale 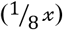. Therefore, the confidence score in this scale has been chosen to detect the existence of the needle more reliably. Let 𝓌^2^, 𝓌^4^ and 𝓌^8^ be the spatial windows of size 4 × 4, 2 × 2 and 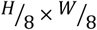 respectively. Firstly all the positions with the confidence score of above *c*_*th* in the whole spatial positions of tensor 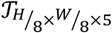 is extracted in 𝕃^8^. In the second step, all the line candidate’s scores are sorted and processes in decreasing order of confidence score in 𝕃^8^.

The selected line 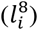 is considered to match with all lines in spatial neighborhood of 𝓌^2^ and 𝓌^4^ in 𝒯^2^ and 𝒯^4^respectively. Finally, all the matched lines are averaged to get a more precise and noise-free estimation of the needle shaft. This procedure is represented in Algorithm 1 in more detail. We chose *c*_*th*_ = 0.9 to select only the line candidate with a confidence score of above 90 percent. *m*_*th* and *θ*_*th*a re the specified threshold value for line matching. These procedures are defined by “*line_candidate*” and “*matching*” in Algorithm 1 respectively.

#### Algorithm 1 *Inference*: outlier removal and hierarchical line estimation

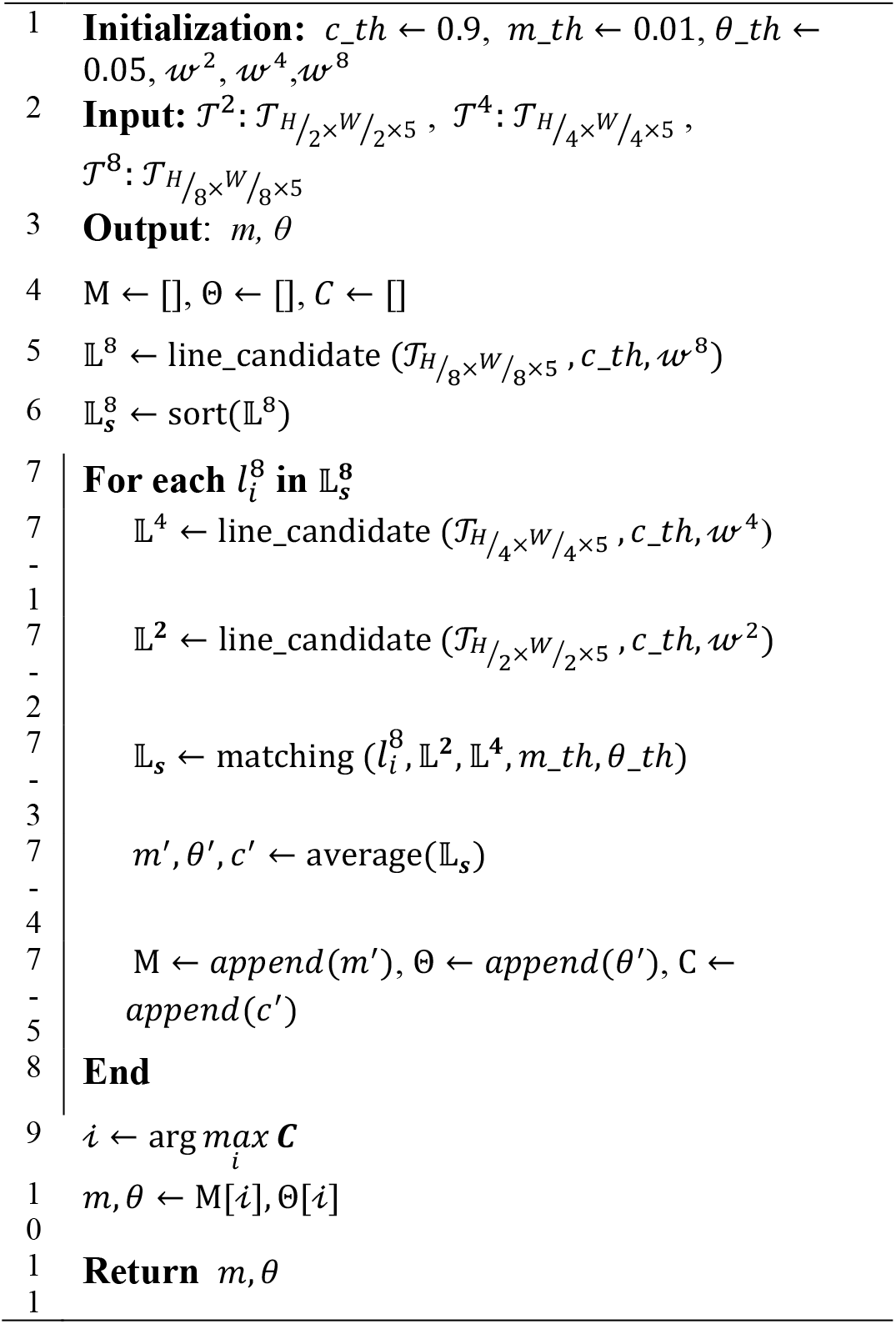

### E. Dataset Overview

US images were captured using a Telemed ultrasound imaging system with a L12 linear array and C6 curvilinear transducers for this study. The dataset was obtained using two phantoms, Ultrasound Compatible Lumbar Epidural Simulator (Kyoto Kagaki co) and femoral vascular access ezono test phantom. A 17 gauge Tuohy epidural needle and a 23-gauge injection needle were utilized to gather in-plane and at angles ranging from 40 to 70 degrees from the body surface normal and from both sides of the transducer. To collect more diverse training data, Needle injections are done in a range from perfectly aligned to 1 cm of normal displacement from the US plane. This method enables the network to train on both visible and invisible needles.

We tracked both the US probe and the needle using an electromagnetic tracker as a gold standard for the shaft and tip localization. In addition, we employed the plus toolkit [65] to temporally calibrate the US frames and determine the transformation between the probe body and the needle base and tip in the reference coordinate system. A total of 10,000 US frames were acquired from 14 videos, comprising seven sequences from a vascular access ezono test phantom and seven sequences from a lumbar epidural phantom using linear and curvilinear transducers.

Three radiologist with more than 4 years of expertise analyzed the labeled dataset to split the dataset for evaluation purposes. We divided the total dataset into two categories, Simple and Complex. These sub-categories are described in further detail below,

- **Simple (C1):** In this category, the needle is aligned correctly with the US plane and therefore emerges as a line-like high-intensity pixels that can be identified statically.
- **Complex(C2):** In this category, the needle is further away from the US plane, making it undetectable to the naked eye. It appeared as moving speckles with slight intensity variation and significant frequency, imperceptible to the naked eye.

## III. Experimental results

The base network (nYolo) was composed of two main components, MFE and AE modules. We performed a comparative evaluation of our proposed method using multiple experiments to show the impact of each model component on the overall performance. In section *A*, we examine the performance of the proposed resolution-preserving network compared to the encoder-decoder version of that called U-Net. In section *B*, we qualitatively investigate the effectiveness of the AE on the selected feature map. Finally, in our last experiment in section *C*, we discuss and show the overall performance of the model components by incorporating them into our proposed nYolo network.

### A. Motion Field Estimation (MFE)

We compared the performance of the proposed RMF with the encoder-decoder version of that given in Balakrishnan’s study[44]. The anticipated motion field was applied to the needle mask in the moving image, and the Dice score was calculated between the needle mask in the fixed image and its corresponding mask in the moved image. As seen in Table I, the proposed RMF when compared with state-of-the-art U-Net, and while the prediction was made only at a fraction of the U-Net parameters, it produces slightly improved accuracy in terms of dice coefficient. The network should have a large enough receptive field and deep features to capture objects and their movements at multiple scales. In the RMF network, the feature tensors are extracted in a hierarchical way using dilated filters. These feature maps are concatenated at the final layer. This structure is beneficial for incorporating spatiotemporal features from very small to larger objects (and their movements) in the final detection layers.

**TABLE I.** The result of the deformable alignment of two consecutive needle masks in US frames using the predicted deformable motion field. The performance is evaluated in terms of average Dice score and runtime. Standard deviation across runs in 5-fold cross validation is included in parenthesis. Our network yields comparable results to U-Net in terms of dice score while operating faster in the inference stage using fewer parameters.

| *Model* | *Dice (% ± std)*<br>Needle Shaft | *GPU (ms)* |
| --- | --- | --- |
| U-Net | 83.5 ± 1.45 | 46 |
| Proposed MFE (RMF) | 84.1 ± 1.21 | 11 |

Fig. 6 shows some examples of the predicted motion field. For the sake of visualization, we set *λ* = 0.05 (3) in Fig. 6 and 0.005 in the final system. Increasing *λ* will result in a smoother motion field, which is not ideal in the final system. The system should keep track of minor changes in the speckles, so that the spatiotemporal features can be used for dynamic detection in the base network.

**Fig. 6.**
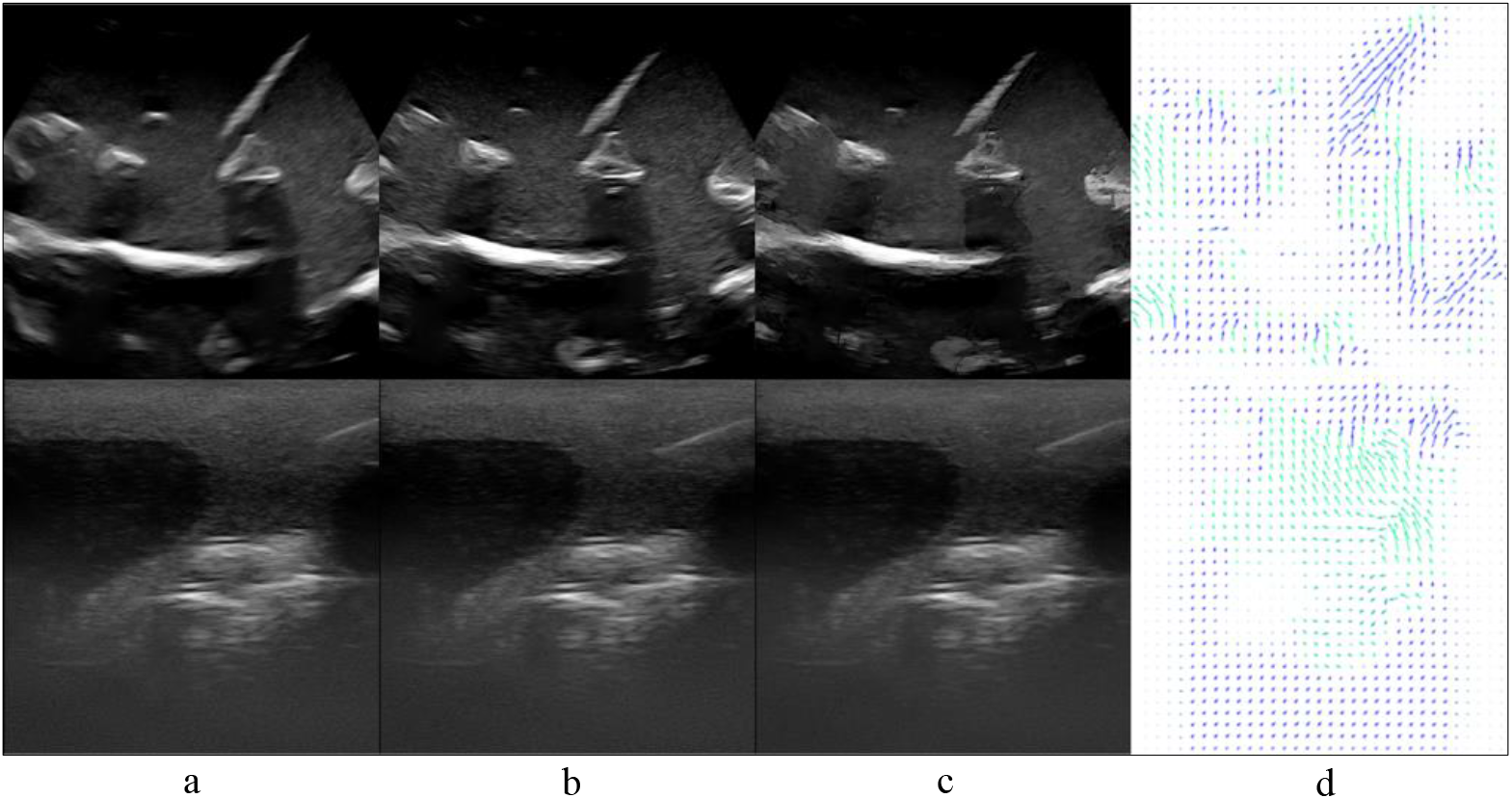
Some examples of the predicted motion field using an unsupervised motion field estimation network (RMF), a) moving, b) fixed, c) moved and d) motion field vectors. We set *λ* = 0.05 (3) for the sake of visualization. With this value, the motion field vector becomes smoother toward prominent variation in the US frame.

The training dataset was generated by concatenating consecutive frames with randomly selected strides of 1 to 5. The dataset is split into three portions for training, validation, and testing: 70%, 15%, and 15% of the whole dataset, respectively. The network was trained for 200 epochs and using the Adam optimizer with a learning rate of 0.001. The weights from the best validation metric were saved for inference during training.

### B. Attention mechanism

Fig. 7 represents some examples of applying the AE module on the feature maps. To evaluate the impact of AE on the feature maps, we chose samples from both the C1 and C2 sub-categories. According to Fig. 7, the first two samples are selected from set C1 and samples 3 to 6 from set C2. US frames have line-like salient intensities in C1. In contrast, the needle characteristics appeared as spatiotemporal features in C2 and must be distinguished using dynamic analysis. As seen in these cases, regardless of the invisibility of the needle in US frames, the feature maps revealed distinct patterns. However, these features have a lot of significant activations in the background which could degrade the Specificity accuracy. Column (c) shows the output maps after applying the AE to the feature maps from the selected layers. As can be observed, this mechanism reduces background noise while highlighting foreground activations. This technique has the potential to improve the training for hard negatives more than for easy negatives and reduce the number of false positive predictions. The last column in Fig. 7 depicts the ground truth bounding boxes that are used in the CL strategy. Because these ground truth boxes are not available during the inference stage, the excitation power is reduced until the end of training to adapt the network for inference.

**Fig. 7.**
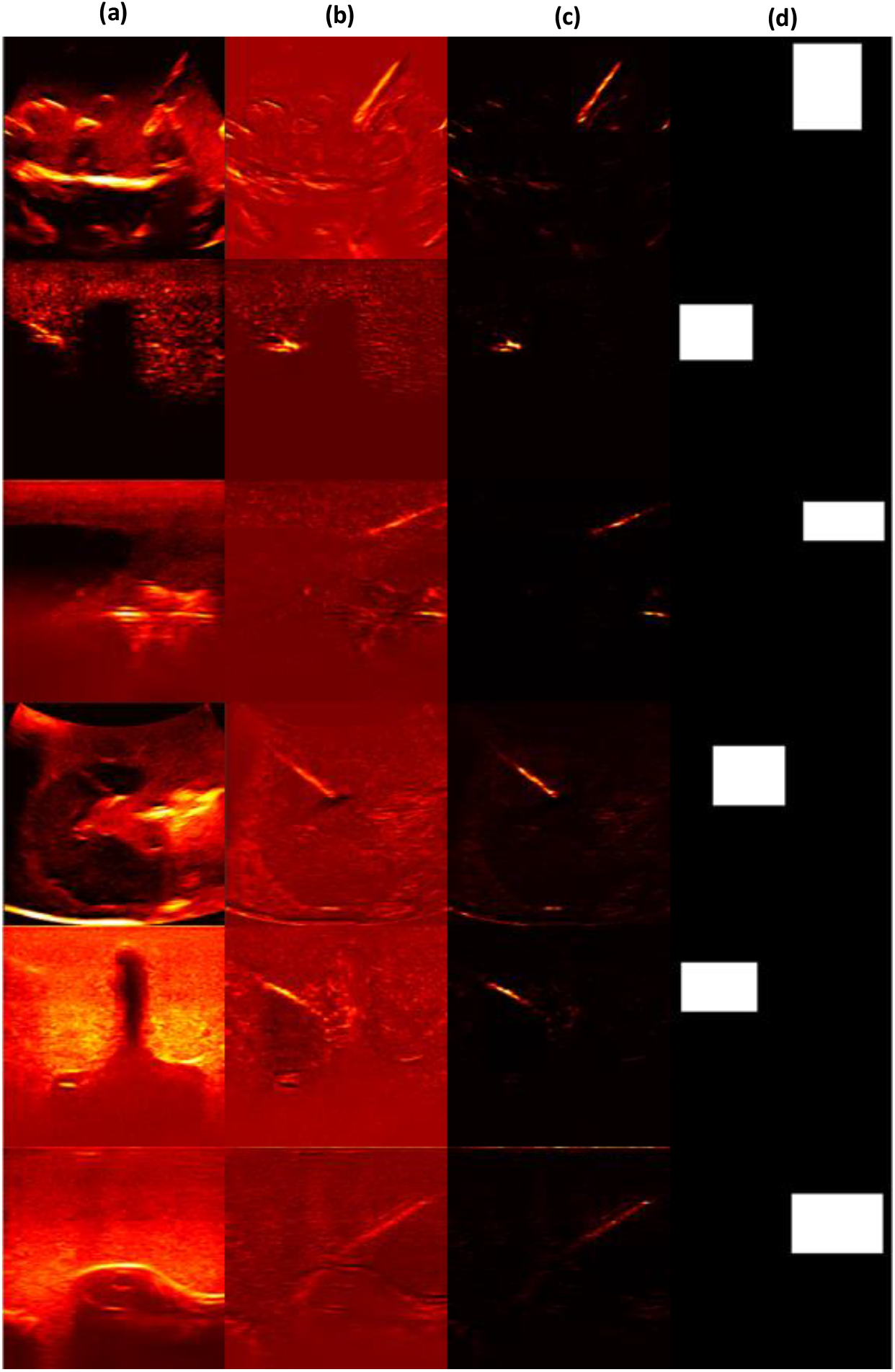
Assisted excitation (AE) to focus on the target regions, a) 2D US frames, b) the feature maps chosen from the neck section’s final layer, c) the resulting attended feature maps using AE, and d) the ground truth bounding boxes

Here, thorough ablation studies are performed to analyze the contribution and effect of several excitation function settings on the *Sensitivity* and *Specificity* of the predictions in our proposed model. We performed six independent experiments to examine the effectiveness of stop_curr variations. Also, in the last experiment we investigated the potential of our proposed Adaptive Excitation scenario. As shown in Fig. 8, the final accuracy degrades significantly in experiment 6. In this experiment, stop_curr was set to 60 and therefore the network was not sufficiently adapted to the inference phase.

**Fig. 8.**
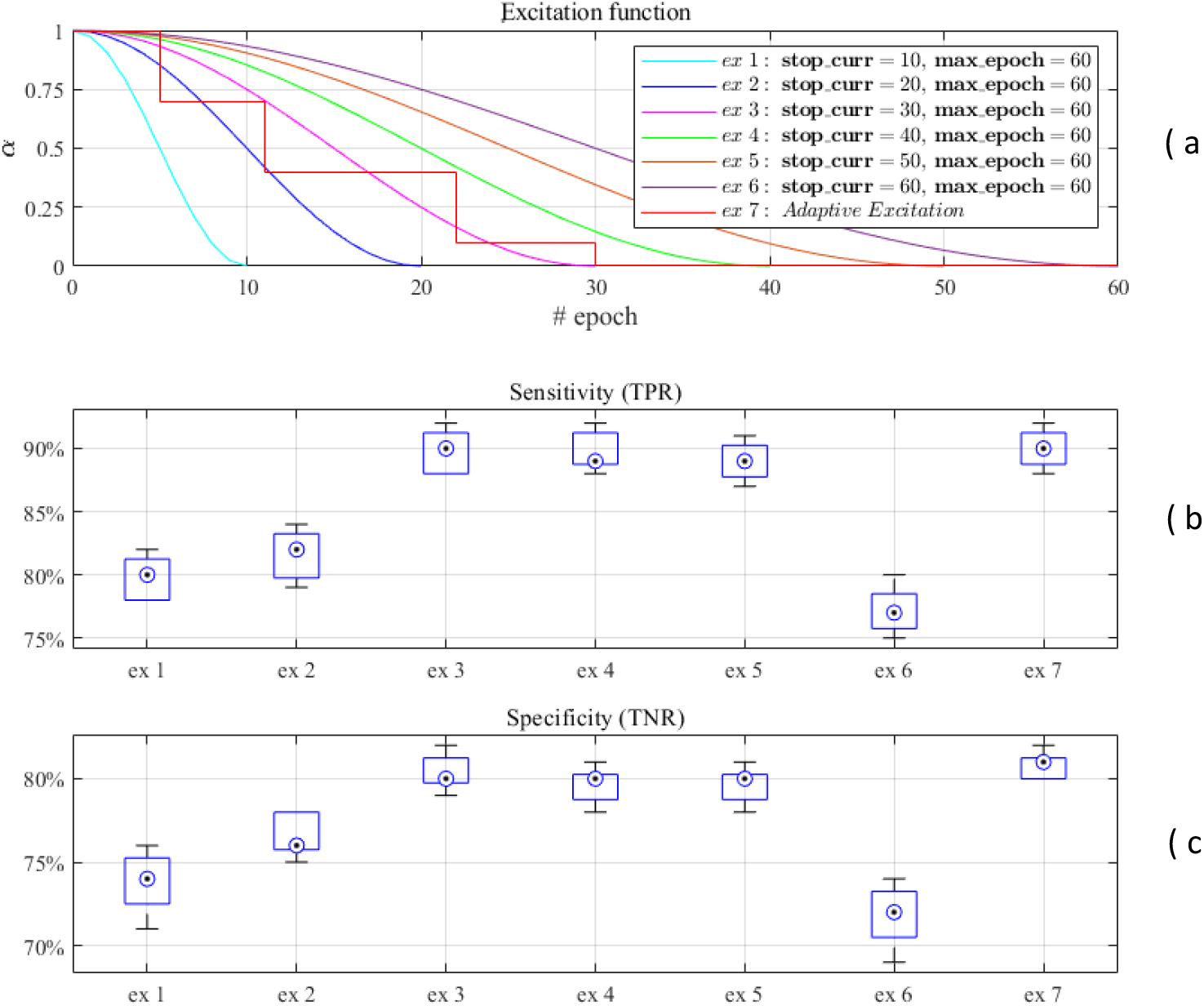
The effect of various excitation scenarios on the accuracy in terms of *Sensitivity* and *Specificity*, a) Seven experiments have been performed on the excitation function with various *stop*_*curr*. The definition of excitation function is given in (8). We chose *max*_*epoch* = 60 empirically for all 7 experiments. b) and c) Statistical analysis of the accuracy in terms of *Sensitivity* and *Specificity*. The minimum accuracy was shown in Experiment 6, where *stop*_*curr* = 60 was selected. In this experiment, the network has no time to adapt itself for the inference. We get the best accuracy in experiments 3 and 7. Using adaptive excitation in experiment 7, the network can automatically adapt itself to the best accuracy in validation without trial and error.

Hence, the rate of false positives and false negatives have increased, which reduces the final accuracy in terms of *Sensitivity* and *Specificity*. Fig. 8 (b, c) also shows that the best results were obtained by experiments 3 and 7. Moreover, the proposed scenario automates the CL process using an adaptive update during training. So that, after several epochs without improvement in the validation metric, the excitation factor decreases by 0.3.

### C. nYolo, needle detection network

We evaluated the performance of our method by comparing the results of the proposed nYolo network with the state-of-the-art Yolov3 framework. To adapt the Yolov3 to our intended application, several ideas from the literature have been brought together and presented in the proposed nYolo. Detection layers and the losses of the Yolov3 were replaced with those in nYolo to adapt it to our application. Moreover, we included GF, RMF, and AE blocks in the Yolov3 and nYolo to analyze the impact of each component on the final accuracy. Table I summarizes the results for two sub-categories, C1 and C2. The first category (C1) refers to most research where the needle is clearly visible in each frame independently. Needle guidance using the thin plane of US imaging is challenging and the alignment is practically impossible. So the second sub-category (C2), which is the subject of our study, deals with the more complicated and probable states in US guided needle injections. Under these conditions, the needle appears as a dynamic of speckles in spatiotemporal patterns. We studied the effect of the motion model as well as attention to improve needle localization in conditions where the needle is statically entirely invisible. We assessed the findings by measuring the shaft angle and tip localization error statistics. The results are summarized in Table II.

**TABLE II.** Ablation study on the effect of each component on the overall accuracy of the proposed needle shaft and tip localization network compared to state-of-the-art Yolov3. The results are given in terms of Sensitivity (TPR) and Specificity (TNR), and the mean angle and tip localization error. The variance of the prediction are measured using standard deviation (Σ) and root mean square error (RMS).

| *Method* | *Sub-category* | *Success* | $\bar{\Delta\theta}$ | $\sigma_{\Delta\theta}$ | $RMS(\Delta\theta)$ | $\bar{\Delta p}$ | $\sigma_{\Delta p}$ | $RMS(\Delta p)$ |
| --- | --- | --- | --- | --- | --- | --- | --- | --- |
| Yolov3 | *C1* | TPR: 100%<br>TNR: 76% | 1.81° | 0.89° | 1.03° | 1.7 | 0.8 | 0.9 |
|  | *C2* | TPR: 62%<br>TNR: 53% | 10.03° | 9.23° | 11.03° | 6.3 | 4.11 | 5.23 |
| Yolov3+GF | *C1* | TPR: 100%<br>TNR: 76% | 1.75° | 0.78° | 0.95° | **1.15** | 0.73 | 1.09 |
|  | *C2* | TPR: 71%<br>TNR: 65% | 3.44° | 2.02° | 2.87° | 3.3 | 2.11 | 2.37 |
| Yolov3+RMF | *C1* | TPR:100%<br>TNR: 79% | 1.53° | 0.9° | 1.12° | 1.19 | 0.7 | 1.06 |
|  | *C2* | TPR: 72%<br>TNR: 69% | 3.05° | 1.87° | 2.24° | 3.4 | 1.75 | 2.08 |
| Yolov3+ RMF +AE | *C1* | TPR:100%<br>TNR: 81% | 1.5° | 0.43° | 1.03° | **0.7** | 0.43 | 0.65 |
|  | *C2* | TPR: **74%**<br>TNR: **71%** | 2.43° | 1.14° | 1.55° | 2.3 | 1.76 | 1.88 |
| nYolo | *C1* | TPR:100%<br>TNR: 78% | 1.75° | 0.83° | 1.08° | 1.54 | 0.74 | 0.83 |
|  | *C2* | TPR: 70%<br>TNR: 63% | 9.45° | 7.21 | 8.67° | 6.52 | 5.33 | 6.04 |
| nYolo +GF | *C1* | TPR:100%<br>TNR: 79% | 1.43° | 0.75° | 0.86° | 1.15 | 0.65 | 0.83 |
|  | *C2* | TPR: 75%<br>TNR: 71% | 3.26° | 2.09° | 2.49 | 3.21 | 2.1 | 2.09 |
| nYolo + RMF | *C1* | TPR:100%<br>TNR: 81% | 1.35° | 1.02° | 1.33° | 1.2 | 0.66 | 0.83 |
|  | *C2* | TPR: 75%<br>TNR:72% | 3.02° | 1.35° | 2.1° | 3.3 | 1.21 | 1.93 |
| nYolo + RMF +AE | *C1* | TPR:100%<br>TNR: 84% | **1.28°** | 0.23° | 0.73° | **0.7** | 0.4 | 0.66 |
|  | *C2* | TPR: **79%**<br>TNR: **74%** | **2.08°** | 1.18° | 1.2° | **2.12** | 1.43 | 2.65 |

The proposed nYolo cuts down the number of parameters in Yolov3 by up to 98% while still providing superior detection accuracy. Indeed, Yolov3 is designed for large-scale datasets, including PASCAL VOC [66] and COCO [67], which have more than 330 thousand images and 80 classes, while in our study, there are only two classes of needle shaft and corresponding tip.

Furthermore, instead of the 9 anchor boxes in each grid cell, the regression network should determine line parameters in our application. We built our framework using these factors. As seen in Table II, the Specificity of the final predictions on C1 has improved from 76% to 78% using Yolov3 and nYolo, respectively. These enhancements are particularly noticeable in C2, where the *Sensitivity* and *Specificity* are boost by 8% and 10% using the two networks. Therefore, considering this significant improvement, it can be confirmed that Yolov3 with about 65 million parameters, which causes overfitting in training, will reduce the accuracy of the inference stage. It is worth noting that the *Sensitivity* of the two networks is 100% on the C1. All of the positive samples are correctly predicted in the C1 sub-category, while many false positives are predicted that have reduced the Specificity. Significant improvements are achieved by incorporating the motion field to the current detectors.

The Gunnar-Farneback (GF) optical flow and the RMF techniques are employed to add extra motion parameters into the system. This method had significant improvements in the *Sensitivity* and *specificity* on the C2, in which the accuracy was enhanced by 9% and 12%, respectively. The discriminative characteristics emerge as moving speckles in multiscale settings in C2. Aside from the significant improvement when employing the motion field, the RMF performs better in terms of inference time and accuracy. In the nYolo base network, the best accuracy was reached by combining RMF with AE. The nYolo network with embedded RMF and AE modules, has a *Sensitivity* and *Specificity* of 79% and 74%, respectively, which is 5% and 3% higher than the Yolov3+RMF+AE framework.

Yolov3 and nYolo have almost 10 degrees and 6 millimeters of error in shaft and tip localization in C2. The addition of motion file estimation blocks, GF or RMF blocks have made significant improvements. Assisted excitation (AE) improves localization and ignores easy negatives.

So it has a significant effect on Specificity in which many of the false alarm outputs disappear. In terms of mean angle error, although we have an acceptable improvement in C1, a significant improvement of this parameter in C2 has been achieved using GF. As seen in Table II, the mean angle error is more than 6 degrees in Yolov3 and nYolo experiments on the C2. Based on these experiments, the minimum error has occurred using nYolo with RMF and AE. The mean angle and tip localization are 2.08 ± 1.18° and 2.12 ± 1.43 mm using nYolo in C2, slightly better than the same configuration using Yolov3.

The software processing unit must operate in real-time to make the system usable in clinical setup. We study the final framework’s performance by including the proposed components in terms of the number of network parameters, processing time and Frame Per Second (FPS) in Table III. The proposed nYolo has less than 1.8% of the total number of Yolov3 weight parameters, and therefore less GPU processing time. We ran the GF optical flow on the CPU, and it took almost 70 milliseconds for every pair of consecutive frames. We proposed the RMF, a motion field estimation network using resolution preserving dilated filters. The proposed network has almost 200 thousand of parameters, and could be processed on GPU by adding less than 3 milliseconds of the processing on the total time of the nYolo.

**TABLE III.** The real-time capability of the proposed nYolo networks against the Yolov3 by embedding the proposed techniques (GF, RMF). The AE module only affects the training process and does not influence the inference stage. therefore, this technique is not mentioned in this table. We used the number of the network parameters, processing time and Frame Per Second (FPS) to compare each setting’s performance in the clinical setup.

| *Method* | *Parameters (M)* | *Time (ms)* | *FPS* |
| --- | --- | --- | --- |
| Yolov3 | 65 | 68 | 14 |
| nYolo | 1.2 | 26 | 38 |
| nYolo+GF | 1.2 | 96 | 10 |
| nYolo+RMF | 1.4 | 29 | 34 |
| nYolo+RMF+Postprocessing | 1.4 | 49 | 20 |

As mentioned earlier, the output prediction tensors should be processed in three scales. We proposed a hierarchical approach from the lowest to the highest scale, to reduce the prediction noise using averaging. This algorithm takes almost 20 milliseconds to get the final needle line parameters and the corresponding tip. So, the final system takes almost 49 milliseconds and the FPS of 20, which is sufficient for real-time operations such as our US-guided needle navigation.

Fig. 9 and Fig. 10 represent qualitative examples of the predicted lines using Yolov3 and the proposed nYolo. The results are depicted on C1 and C2 sub-categories and by embedding the GF, RMF and the AE modules to the base networks. The examples of rows (a) and (b) are selected from the C1 and the others are selected from the C2. Although the needle is more discriminative in the C1, it could be seen in Fig. 9, rows (a) and (b) that utilizing the GF and RMF enhances the shaft and the tip localization using Yolov3 obviously. Further, AE enhances the accuracy in terms of the Specificity, resulting in fewer false positive predictions. This helps the proposed hierarchical post-processing algorithm with less outlier and noise-free predictions in higher scale outputs. This observation is also obvious in Fig. 10.a and Fig. 10.b using nYolo. As can be seen in Fig. 10, these enhancements are more obvious utilizing the GF and RMF in the nYolo framework. This is because the nYolo is designed and optimized with fewer parameters for our application, therefore the network has more capability to confront unseen samples in the inference stage. In general, nYolo has minor error in the shaft and tip localization. This superiority is more evident in C2 sub-category.

**Fig. 9.**
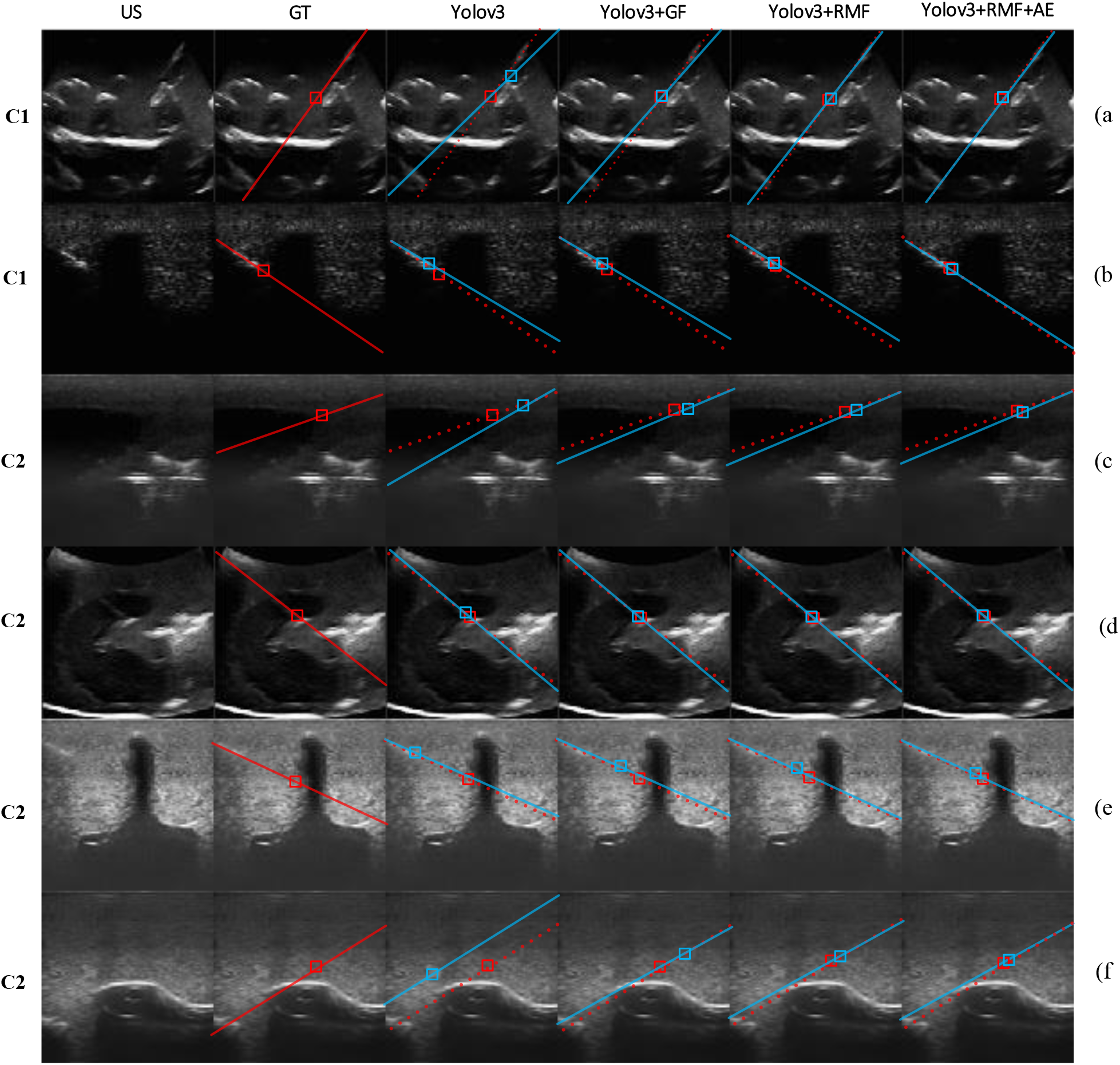
Needle shaft and tip localization in 2D US sequences using Yolov3 architecture. The results are presented on the C1 and C2 sub-categories. The category of each example is depicted in the left column. The best TNR and accuracy are given by utilizing RMF and AE together. As mentioned, AE besides handling class imbalances, also gives more attention to hard negatives and therefore the accuracy has improved. The red line represents the ground truth and the prediction has depicted in blue.

**Fig. 10.**
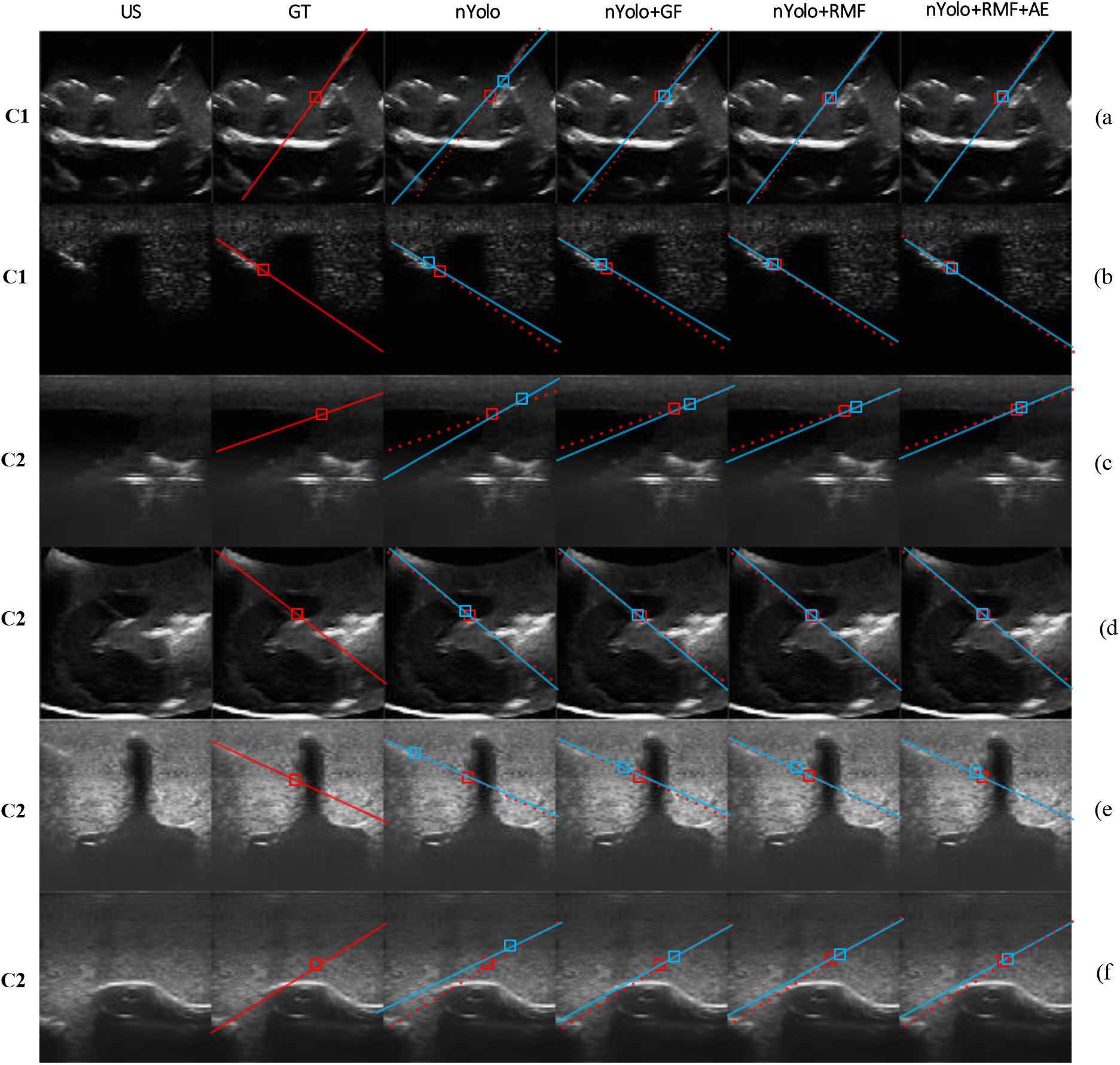
Needle shaft and tip localization in 2D US sequences using the nYolo architecture. The proposed architecture utilizes residual and ASPP blocks. The results show that the localization is less biased than Yolov3 architecture (Fig. 9). The red line represents the ground truth and the prediction has depicted in blue.

The needle is poorly visible in the C2 compared to the C1 sub-category. In this situation, the prominent discriminative features of the needle are highlighted using dynamic analysis of the speckles. The examples in rows (c) to (f) in Fig. 9 and Fig. 10 are related to the C2. For example, It can be seen that the needle is completely invisible in rows (c) and (f), and therefore the predicted line is vulnerable to noise and outlier in the predictions. Further, the final line angle error in row (c) is more than 10 degrees without proposed components. As seen in column 3, embedding GF improves the angle error and the tip localization, respectively. In addition to being significantly effective, this powerful technique suffers from substantial CPU processing time, which degrades the real-time capability of the system in the real clinical settings. Moreover, the RMF is based on CNNs and can be embedded in the Yolov3 or nYolo frameworks. In addition to a slight improvement in overall accuracy compared to GF, this technique has excellent real-time performance, which adds only about 3 milliseconds of GPU processing time to the base networks. As seen in Fig. 9 and Fig. 10, the RMF method has slightly better angle and tip localization capability in nYolo than Yolov3.

The last column in Fig. 9 and Fig. 10 represents the final predictions using the AE and RMF modules within the base networks. AE module improves the predictions by reducing the false positive rates. As a result, the angle of line candidates, and the tip localization are improved moderately. For example, considering row (e), the needle tip is ambiguous. Therefore, the tip is localized with huge error using nYolo and Yolov3 frameworks, although acceptable improvement is achieved using GF and RMF, the tip is localized significantly using AE module.

## IV. Discussions

In an effort to achieve a top-performing needle detection system, we proposed an efficient CNN network and employed several techniques to enhance the accuracy as well as the real-time capability. We gathered and divided the dataset into two sub-categories. In the first sub-category (C1), the needle appears as line-like high intensity regions, which could be recognized statically. This situation is the focus of many studies in the literature. Aligning the needle with the thin plane of the transducer is challenging, which leads to ambiguity in 2D US frames independently. Therefore, the second sub-category (C2) consists of frames in which the needle is invisible to the naked eye. In these situations, the needle can be recognized by analyzing the dynamic characteristics of the speckles captured using a spatiotemporal feature detector. We developed our CNN network based on Yolov3, a state-of-the-art object detection framework. Furthermore, the main feature extractor blocks are selected to extract spatial (and not temporal) features. The Yolov3 architecture has more than 65 million of parameters that could lead to over-fitting in our application. We have a maximum of two classes for needle shaft and tip in one frame. Therefore, this framework could not be beneficial for our application, in which the main discriminative patterns are embedded in the spatiotemporal patterns of speckle dynamics. We proposed a modified version of Yolov3 named nYolo. In this framework, we utilized two techniques in the base network to improve the accuracy, the Resolution Preserving Motion Field estimation network (RMF) and the CL-based attention mechanism. We extract the motion field using RMF and augment the training frame sequences with the amplitude and phase of the field. Our newly developed RMF performs better than U-Net while using far fewer parameters.

Furthermore, the AE module is utilized to improve the accuracy in terms of Specificity by highlighting foreground activations. According our ablation studies on the effect of AE we found that utilizing the CL approach has a considerable impact on the overall accuracy of the line parameter predictions. The key element in the CL approach is the excitation function’s definition (8). We examined the impact of the AE module on feature maps with more prominent distinguishing features for needle detection. We found that many background tissue structures can have a negative impact on the prediction. The activations inside the needle boxes are excited, and therefore the background region is attenuated during training. Using this technique, the network learns to focus more on important regions. After the training, these excitations gradually fade and become zero. The bounding box masks are not accessible at the inference stage. As a result, the network should be trained to make predictions since it has no prior knowledge about the valid needle boxes. The curriculum technique is used to overcome this problem, which involves minimizing the effect of AE on an adaptive update of the excitation factor.

In table II we provide a summary of the analysis performed in this work and highlights the study’s main contributions. Based on quantitative and qualitative evaluations, we found that the proposed approach was achieve significant improvement by adding the RMF and AE to the base network in the C2 sub-category. The overall performance of the final system (nYolo+RMF+AE) is enhanced compared to the Yolov3+RMF+AE by 0.35° and 0.18 millimeters in terms of mean angle and tip localization error, respectively.

The real-time capability of the system is also investigated. As seen in Table III, the proposed framework has almost 1.4 million of parameters and comes with an end-to-end one-stage mechanism. As a result, considering the post-processing stage, the system could operate at 20 FPS. Overall, the proposed system can achieve an adequate performance in US-guided needle detection scenarios. Since the framework is designed to capture both the static and dynamic patterns, the system could operate reliably without missing the tracking during the needle insertion procedures.

## V. Conclusions

In the present work a lightweight needle detection framework based on speckle motion analysis was designed, implemented, and tested. In this study, we bring together several ideas from the literature and present them in a unified end-to-end one stage needle detection network. We explored our proposed model thoroughly in US-guided needle navigation systems and in scenarios where the shaft and the corresponding tip are ambiguous. From our experiments, we concluded that analyzing the dynamicity of the speckles has enough discriminative characteristics to overcome the challenging problem of needle tracking in real-world clinical routines. In addition, the real-time capability of the system was confirmed to offer a clinically accepted mechanism. Our experiments demonstrate the effectiveness of the proposed needle detection framework. This study provides helpful insights into future deep learning-based works on needle-like tool detection in ultrasound interventions for the first time in the literature.

